# Circadian key component CLOCK/BMAL1 interferes with segmentation clock in mouse embryonic organoids

**DOI:** 10.1101/2020.10.30.362830

**Authors:** Yasuhiro Umemura, Nobuya Koike, Yoshiki Tsuchiya, Hitomi Watanabe, Gen Kondoh, Ryoichiro Kageyama, Kazuhiro Yagita

## Abstract

In mammals, circadian clocks are strictly suppressed during early embryonic stages as well as pluripotent stem cells, by the lack of CLOCK/BMAL1 mediated circadian feedback loops. During ontogenesis, the innate circadian clocks emerge gradually at a late developmental stage, then, with which the circadian temporal order is invested in each cell level throughout a body. Meanwhile, in the early developmental stage, a segmented body plan is essential for an intact developmental process and somitogenesis is controlled by another cell-autonomous oscillator, the segmentation clock, in the posterior presomitic mesoderm (PSM). In the present study, focusing upon the interaction between circadian key components and the segmentation clock, we investigated the effect of the CLOCK/BMAL1 on the segmentation clock *Hes7* oscillation, revealing that the expression of functional CLOCK/BMAL1 severely interferes with the ultradian rhythm of segmentation clock in induced PSM and gastruloids. RNA sequencing analysis showed that the premature expression of CLOCK/BMAL1 affects the *Hes7* transcription and its regulatory pathways. These results suggest that the suppression of CLOCK/BMAL1-mediated transcriptional regulation during the somitogenesis may be inevitable for intact mammalian development.

## Introduction

The circadian clock is the cell-autonomous time-keeping system generating the orderly regulated various physiological functions, which enables cells, organs, and systems to adapt to the cyclic environment of the rotating Earth (1-5). The core architecture of the circadian molecular clock consists of negative transcriptional/translational feedback loops (TTFLs) composed of a set of circadian clock genes, including *Bmal1, Clock, Period* (*Per1, 2, 3*), and *Cryptochrome* (*Cry1, 2*), functioning under the control of E-box elements (2, 6). The kernel of TTFLs is composed of heterodimerized CLOCK/BMAL1 key transcriptional factors that positively regulate the circadian output genes, as well as *Per* and *Cry* genes via E-box. PERs and CRYs inhibit CLOCK/BMAL1 transcriptional activity, and the negative feedback loops between these genes generate oscillations of approximately 24 hr.

In mammalian development, it has been demonstrated that early embryos and pluripotent stem cells have no apparent circadian molecular oscillations (7-11), whereas the innate circadian clock develops during ontogenesis and is established at a late developmental stage (12-15). Regarding the mechanisms regulating circadian clock development, using an *in vitro* model of embryonic stem cell (ESC) differentiation and mouse embryos, it was shown that prolonged posttranscriptional mechanisms, such as suppressed translation of CLOCK protein and predominant cytoplasmic localization of PER proteins, inhibit the establishment of the circadian TTFL cycle (16-18). Although it was revealed that the multiple mechanisms strictly suppress the circadian molecular clock in the undifferentiated cells and early stage embryos, the biological and physiological significance of the delayed emergence of circadian clock oscillation in mammalian embryos has been unknown.

In the early developmental stages, a segmented body plan is essential for an intact developmental process. Somitogenesis is related to another cell-autonomous oscillator, the segmentation clock, in the posterior presomitic mesoderm (PSM) (19, 20). The mouse segmentation clock is underlain by a negative feedback loop involving *Hes7* oscillation (21, 22). HES7 is a key transcriptional factor that represses its own expression and oscillates through a negative feedback loop in a period of 2–3 hr in mouse and 4–5 hr in humans. The NOTCH, WNT, and fibroblast growth factor signaling pathways are involved in the regulation of the *Hes7* oscillator and its intercellular synchronization (20). In mammals, two different types of rhythm sequentially emerge during the developmental process, however, there is a lack of knowledge about the biological significance of the rhythm conversion during development.

In this study, focusing on the relationship between circadian clock and segmentation clock, we investigated the effect of the premature expression of CLOCK/BMAL1 on the segmentation clock oscillation, and revealed severe interference with the ultradian rhythm of segmentation clock in iPSM and gastruloids. RNA sequencing analysis showed that CLOCK/BMAL1 affects the *Hes7* transcription and its regulatory pathways. These findings highlight that the suppression of functional CLOCK/BMAL1, which leads to arrest the circadian clock oscillation, during the early to mid-developmental stage may be inevitable for the intact process of mammalian embryogenesis.

## Results

### A circadian clock gene, *Per1*, and a segmentation clock gene, *Hes7*, are adjacent genes in the mammalian genome

In mammals, the temporal relationship between the segmentation clock and circadian clock appears to be mutually exclusive (**Fig. S1**) (12-15, 19, 20). To explore the functional interaction between these two biological rhythms with different frequencies during the developmental process, we focused on the genomic architecture of genes comprising circadian and segmentation clocks, respectively. Intriguingly, one of the core circadian clock genes, *Per1*, is physically adjacent to an essential component of the segmentation clock, *Hes7*, in a genomic region conserved in higher vertebrates, including mice and humans. The *Per1* homolog *Per2* is adjacent to the *Hes7* homolog *Hes6* in the genome (**Fig. 1A**). Since *Hes7* exhibits the essential characteristics of a segmentation clock (23) and neighboring genes can influence the expressions with each other during somitogenesis in zebrafish (24), we focused on the effect of the regulation mechanism of the circadian clock on segmentation clock oscillation. Therefore, we investigated the effect of the CLOCK/BMAL1-mediated activation of *Per1* transcription on the segmentation clock oscillation in induced presomitic mesoderm (iPSM), an *in vitro* recapitulating model of a segmentation clock, using ESCs carrying the *Hes7*-promoter-driven luciferase reporter (*pHes7-luc*) (25) (**Fig. 1B**). In the iPSM, the pluripotent markers have not yet been down-regulated sufficiently as previously reported (25), and the iPSM differentiated from *Per2*^*Luc*^ ESCs showed no apparent circadian clock oscillation (**Fig. S2A and B**). In addition, the immunostaining pattern of CLOCK, BMAL1, and PER1 in the iPSMs was quite similar to that in the undifferentiated ESCs (**Fig. S2C**) (16, 17), confirming that the circadian TTFL was not established and circadian clock oscillation was also strictly suppressed in the iPSM by the common inhibitory molecular mechanisms to the undifferentiated ESCs.

**Fig. 1.**
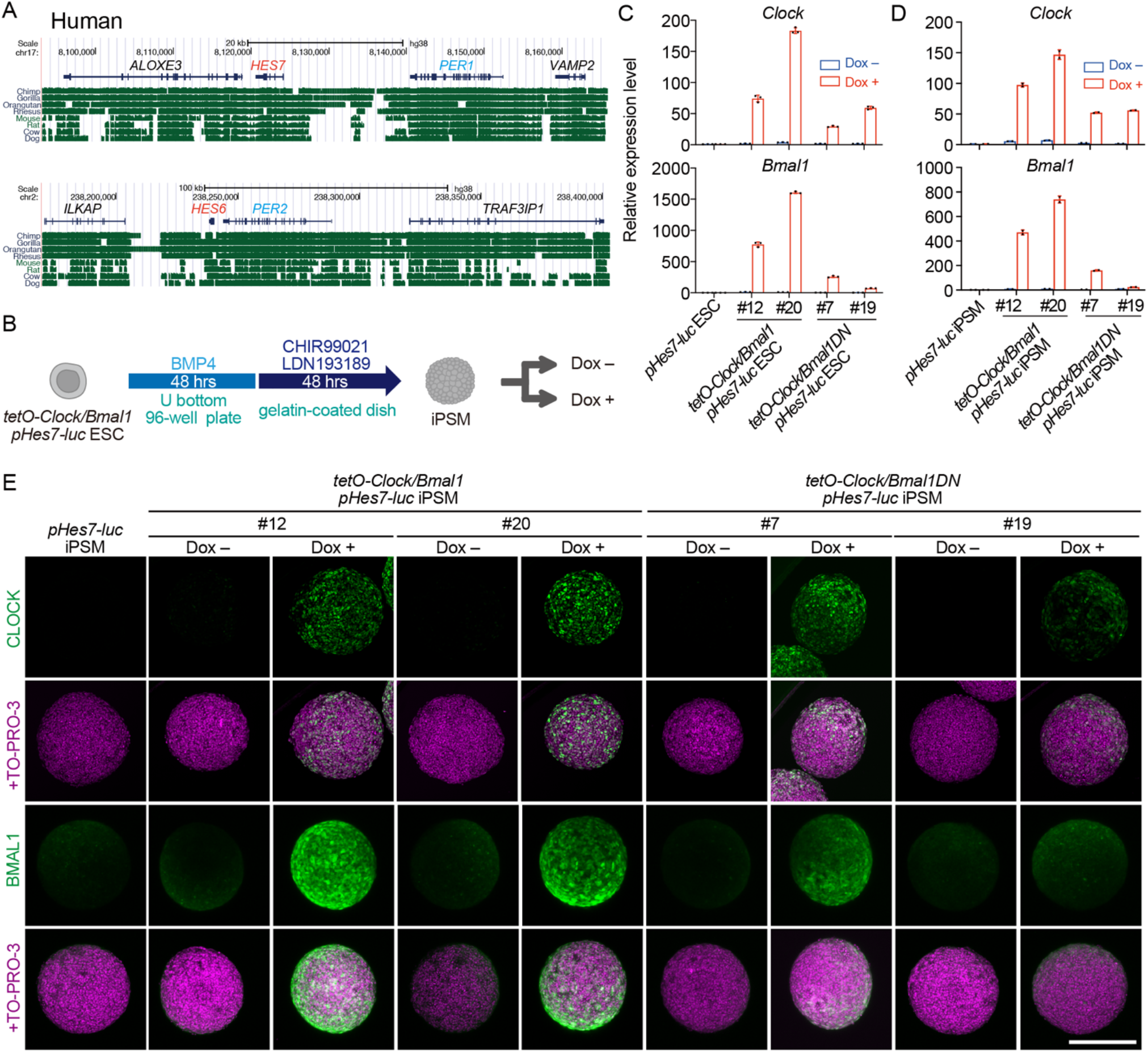
Establishment of ESC lines carrying both the dox-inducible *Clock* and *Bmal1* genes. (**A**) Human genomic locus of circadian clock genes, *PER1*, and the essential segmentation clock gene, *HES7*, are highly conserved in higher vertebrates. The *PER1* homolog *PER2* is also located adjacent to the *HES7* homolog *HES6* in the genome. (**B**) ESCs were differentiated into iPSM for 96 hr *in vitro*, and then the iPSM colonies were treated with or without dox. (**C**) qPCR of *Clock* and *Bmal1* mRNA in the indicated ESCs. 500 ng/mL dox treatment for 6 hr (red) or not (blue). Each number indicates clone number. Mean ± SD (n = 3 biological replicates). (**D**) qPCR of *Clock* and *Bmal1* mRNA in the indicated iPSM colonies. 1000 ng/mL dox treatment for 2 hr (red) or not (blue). Mean ± SD (n = 2 technical replicates). The average expression level of *pHes7-luc* ESCs or iPSM colonies without dox was set to 1. (**E**) Representative maximum intensity projection of the immunostaining of iPSM colonies treated with 1000 ng/mL dox for 2 hr or not. n = 2–3 biological replicates. Scale = 250 µm.

Because CLOCK/BMAL1 is key transcription regulator of circadian TTFL, and the expression of CLOCK protein is suppressed post-transcriptionally in iPSM as well as ESCs and early embryos (17), we established two ESC lines carrying both the doxycycline (dox)-inducible *Clock* and *Bmal1* genes (**Fig. 1C, Fig. S3**). In iPSM differentiated from ESCs, the expression of both *Clock/Bmal1* mRNA and CLOCK/BMAL1 proteins was confirmed after the addition of dox (**Fig. 1 D and E**), and we found that overexpression of both *Clock* and *Bmal1* successfully activated the expression of core clock genes (**Fig. 2A**). As the dominant negative mutant of *Bmal1* (*Bmal1DN*) (26) co-expressed with *Clock* did not activate the *Per1/2* and *Cry1/2* genes, we concluded that CLOCK/BMAL1 specifically activated the expression of these clock genes via an E-box (**Fig. 1 C–E, 2A**). We then examined the expression of genes in *Hes7*, which is proximal to *Per1*, and *Hes6*, which is proximal to *Per2* in the genome. The expression of *Clock/Bmal1* induced by dox in the iPSM induced significant upregulation of the expression of the *Hes7*, but not *Hes6*, gene (**Fig. 2B**). Similarly, we also observed the upregulation of *Hes7* expression by *Clock/Bmal1* induction in the undifferentiated ESCs (**Fig. S4 A and B**). These results indicate that the circadian components CLOCK/BMAL1 also affect the segmentation clock gene *Hes7*, as well as *Per1*.

**Fig. 2.**
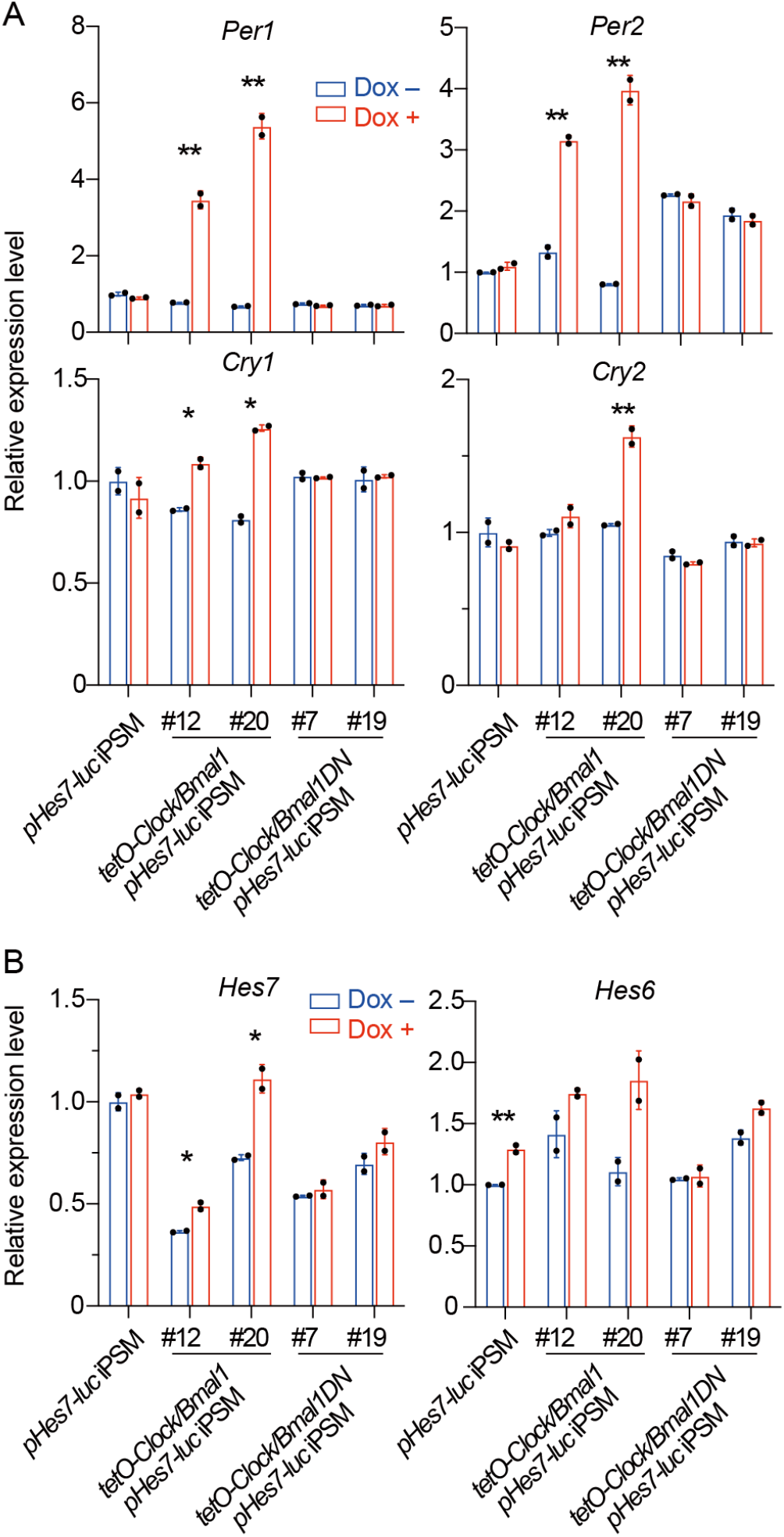
CLOCK/BMAL1 expressions upregulated not only circadian clock genes but also *Hes7* gene expressions in the iPSM. (**A, B**) qPCR of core circadian clock genes (**A**) and *Hes*7 or *Hes6* gene (**B**) in the indicated iPSM colonies. 1000 ng/mL dox treatment for 2 hr (red) or not (blue). Mean ± SD (n = 2 technical replicates). The average expression level of iPSM colonies without dox was set to 1. Two-tailed t-test, *P < 0.05, **P < 0.01.

### Inhibition of *Hes7* ultradian rhythm by CLOCK/BMAL1 in iPSM

We next performed a functional analysis using the *in vitro* recapitulation model of a segmentation clock oscillation in iPSM (25). The oscillations in bioluminescence from *pHes7-luc* reporters were observed using a photomultiplier tube device (PMT) and an EM-CCD camera (**Fig. 3A**). We confirmed an oscillation of *Hes7*-promoter-driven bioluminescence with a period of approximately 2.5–3 hr in control iPSM with or without dox using PMT and the EM-CCD camera (**Fig. 3 B and C**). Traveling waves of *pHes7-luc* bioluminescence were observed, indicating that the segmentation clock oscillation in iPSM was successfully recapitulated, consistent with a previous report (25). Using this iPSM-based segmentation clock system, we investigated the effect of *Clock/Bmal1* expression on *Hes7*-promoter-driven oscillation. The expression of *Clock/Bmal1* genes (Dox+) resulted in defects of the oscillation in *Hes7* promoter activity, whereas *pHes7-luc* bioluminescence continued to oscillate under Dox– conditions. Oscillation of the segmentation clock was observed even during the induction of *Clock/Bmal1DN* (**Fig. 3D**), indicating that the CLOCK/BMAL1-mediated mechanism interfered with the transcriptional oscillation of *Hes7*. A traveling wave of *Hes7* promoter activity disappeared with the expression of *Clock/Bmal1* (**Fig. 3 E and F**), and dox-dependent arrest of *pHes7-luc* traveling wave (**Fig. 3 G and H**) clearly demonstrated the CLOCK/BMAL1-mediated interference with *Hes7*-driven segmentation clock oscillation in iPSM.

**Fig. 3.**
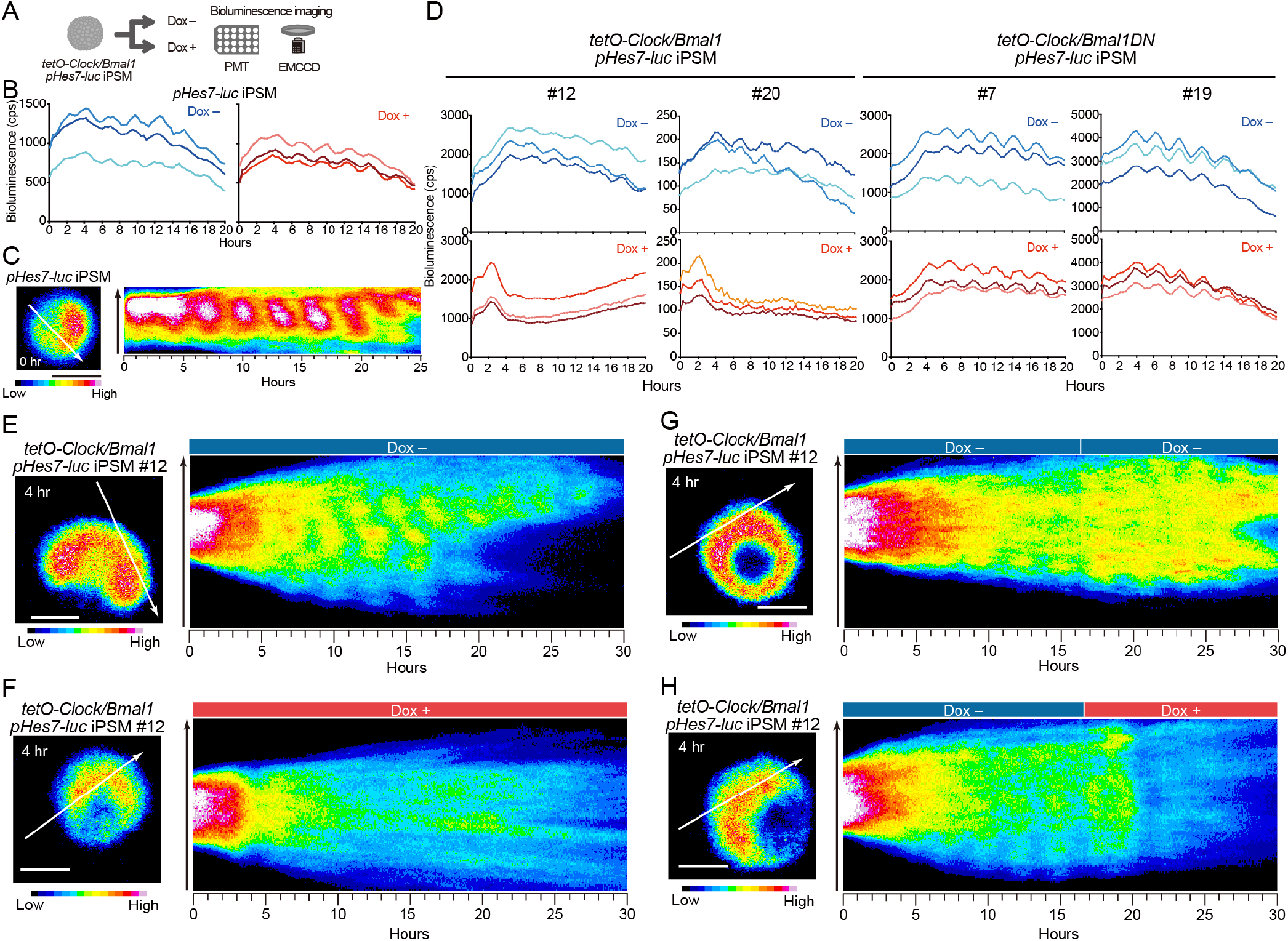
CLOCK/BMAL1 expressions arrested the autonomous oscillations of *Hes7* in the iPSM colonies. (**A**) Bioluminescence of each dox-inducible *Clock/Bmal1* or *Clock/Bmal1DN pHes7-luc* iPSM colony was observed using PMT or an EM-CCD camera without or with dox. (**B, C**) Representative bioluminescence traces (**B**, n = 25 biological replicates) and live imaging (**C**, n = 3 biological replicates) of single *pHes7-luc* iPSM colony with or without dox. The kymograph of the imaging along the arrow is shown. (**D**) Representative bioluminescence traces of the single indicated iPSM colony with and without 1000 ng/mL dox. n = 10-54 biological replicates. (**E–H**) Live imaging of the single *tetO-Clock/Bmal1 pHes7-luc* iPSM colony with and without dox. Dox-containing medium or only medium was added at the indicated time points at the final dox concentration of 1000 ng/mL (**G, H**). Each kymograph along the arrow is shown. Scales = 250 µm. n = 2-4 biological replicates.

### Interference with somitogenesis-like segmentation by induction of CLOCK/BMAL1 in gastruloids

In addition, to explore the effect of CLOCK/BMAL1 expression on somitogenesis, we established the ESC-derived embryonic organoids, gastruloids, recapitulating an embryo-like organization, including somitogenesis-like process *in vitro* (27) (**Fig. 4A**). The *pHes7-luc* bioluminescence represented a traveling wave accompanied by the formation of segment-like structures with anteroposterior polarity, in which the gastruloids were stained with stripes of a somite marker, *Uncx4*.*1*, by *in situ* hybridization (**Fig. 4 B–D**). Only dox treatment in control gastruloids induced no change in the *pHes7-luc* bioluminescence oscillation and somitogenesis-like process (**Fig. 4 E–G**). The dox-inducible *Clock/Bmal1* ESC line carrying *pHes7-luc* was differentiated *in vitro* into gastruloids and produced the somitogenesis-like process without dox (**Fig. 4 H–J**). In contrast, the dox-dependent induction of *Clock/Bmal1* expression in the gastruloids interrupted the *pHes7-luc* oscillation and disrupted the somitogenesis-like structures (**Fig. 4 K–M**). In gastruloids, the expression of both *Clock/Bmal1* mRNA was confirmed after the addition of dox (**Fig. S5**). These results suggest that the premature expression of the circadian key transcriptional regulator CLOCK/BMAL1 critically interferes with not only *Hes7* oscillation, but also somitogenesis.

**Fig. 4.**
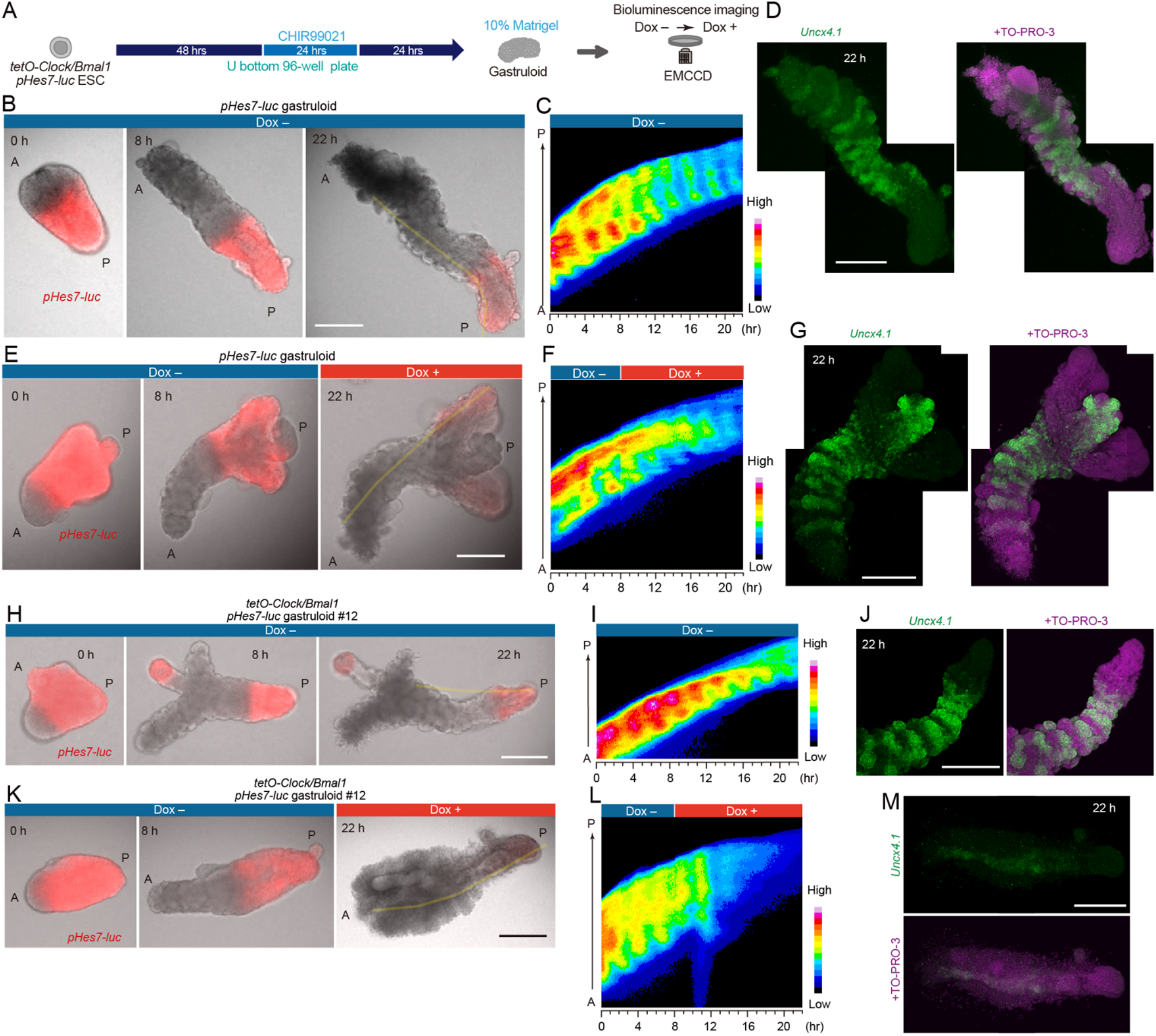
CLOCK/BMAL1 expressions interfered with the autonomous oscillations of *Hes7* and somitogenesis-like process in the gastruloids. (**A**) Dox-inducible *Clock/Bmal1 pHes7-luc* ESCs were differentiated into gastruloids for 96 hr *in vitro*, and then the gastruloids embedded in 10% Matrigel were treated with or without dox. *pHes7-luc* bioluminescence was observed using an EM-CCD camera without or with dox. (**B–M**) Time-lapse bioluminescence (red) and bright field imaging of the single *pHes7-luc* gastruloid or *tetO-Clock/Bmal1 pHes7-luc* gastruloid without and with dox. Dox-containing medium was added at the indicated time points at the final dox concentration of 1000 ng/mL (**C, F, I, L**). Each kymograph is shown along the yellow lines in **B, E, H, K**. *In situ* hybridization of *Uncx4*.*1* in the gastruloids after the live cell imaging (**D, G, J, M**). Scales = 250 µm. n = 2-4 biological replicates.

### CLOCK/BMAL1-mediated interference in *Hes7* regulatory network

Next, to examine the perturbation mechanisms of the segmentation clock oscillation by the circadian components CLOCK/BMAL1, we analyzed the RNA sequencing (RNA-seq) data obtained from the total RNA of iPSM colonies. We extracted 509 upregulated and 88 downregulated differentially expressed genes (DEGs) after the induction of *Clock/Bma11* gene expression in iPSM colonies (**Fig. 5A**). A KEGG pathway enrichment analysis for the DEGs revealed enrichment of the WNT, MAPK, and NOTCH signaling pathways related to *Hes7* oscillation (28) (**Fig. 5B**). Almost all other ranked pathways also included the WNT, MAPK, and NOTCH signaling pathway-related genes (**Fig. 5B**). Similarly, enrichment of the WNT, MAPK, and NOTCH signaling pathways by *Clock/Bmal1* induction was also observed in the undifferentiated ESCs (**Fig. S6 A and B**). These findings indicate that the expression of CLOCK/BMAL1 affects the *Hes7*-related signaling pathways, which interferes with the feedback loop regulating *Hes7* oscillation. Intriguingly, in addition to *Hes7* gene expression, the expressions of *Aloxe3* in iPSM and *Aloxe3* and *Vamp2* in ESCs, the other contiguous genes with *Per1*, were upregulated with the induction of *Clock/Bmal1* expression, and this result was confirmed by quantitative RT-PCR (qPCR) (**Fig. 5 C–E, Fig. S6 C–E**) suggesting that forced expression of CLOCK/BMAL1 also affects a wide region around the *Hes7* gene locus on the same chromosome. These results suggest that the premature expression of the circadian components CLOCK/BMAL1 interfered with *Hes7* oscillation and somitogenesis by perturbing the *Hes7* expressions through indirect regulatory pathways (**Fig. 5F**). Because the loss of the *Hes7* ultradian expression rhythm in the mouse cause segmentation defects (22, 29), the oscillatory expression of *Hes7* is essential for mammalian development. Therefore, the results in this study suggest that it may be imperative that CLOCK/BMAL1 function is suppressed until the completion of segmentation and other related developmental events.

**Fig. 5.**
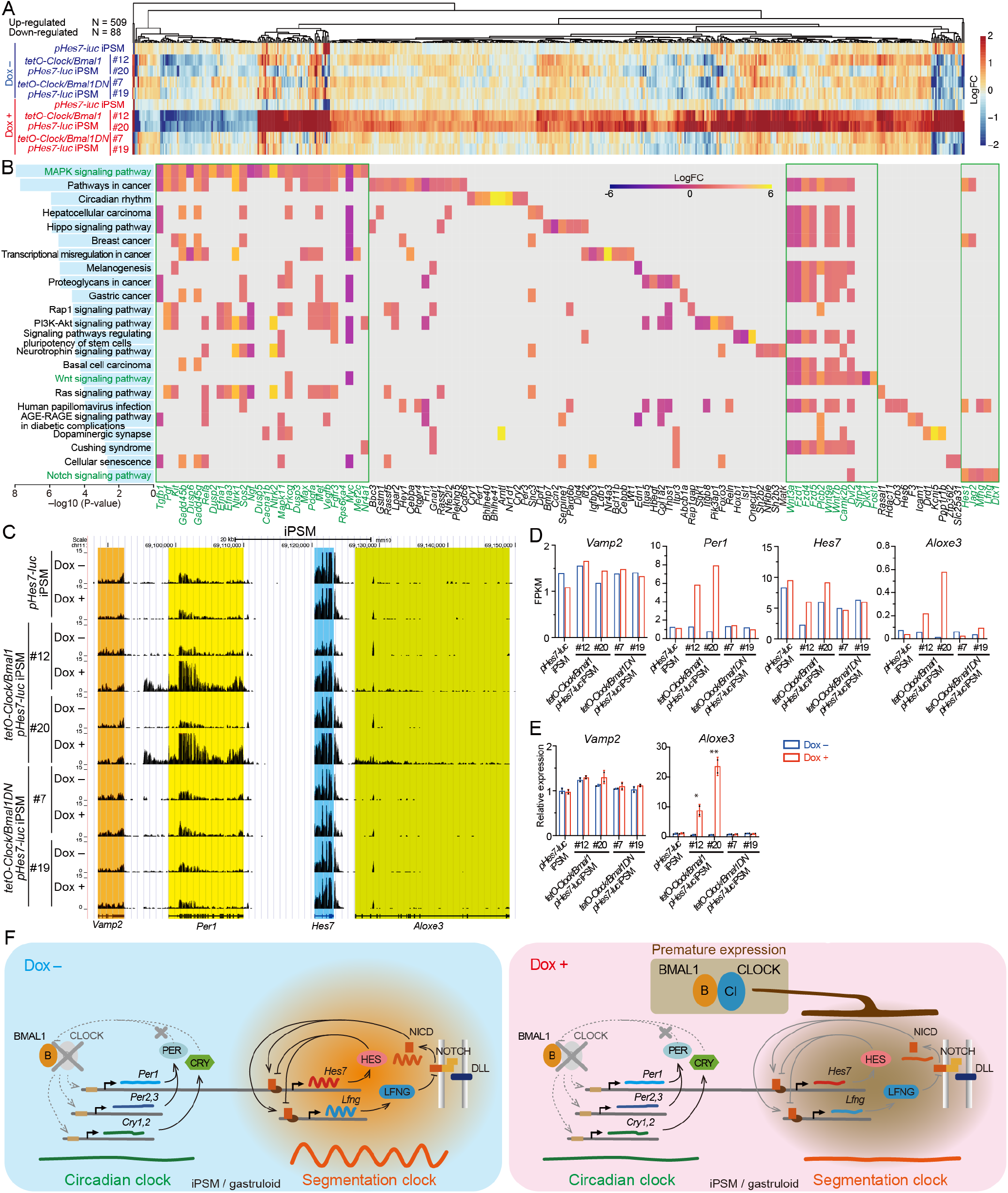
CLOCK/BMAL1 expressions in the iPSM colonies affected *Hes7*-related signaling pathways and upregulated the expression of contiguous genes, *Per1, Hes7*, and *Aloxe3*. (**A**) Upregulated and downregulated DEGs in the indicated iPSM colonies treated with dox. (**B**) KEGG pathway analysis of the DEGs. Each pathway was indicated with each transformed p-value. The ranked pathways contained several common genes in the WNT, MAPK, and NOTCH signaling pathways. (**C**) UCSC genome browser views of RNA-seq data of the contiguous genes *Vamp2, Per1, Hes7*, and *Aloxe3*. The reads shown are normalized average reads per 10 million total reads in 10-bp bins. (**D**) mRNA expression of *Vamp2, Per1, Hes7*, and *Aloxe3* in the indicated iPSM colonies according to RNA-seq. (**E**) Validation of *Vamp2* and *Aloxe3* gene expression levels in the indicated iPSM colonies using qPCR. Colored boxes indicate 1000 ng/mL dox treatment for 2 hr (red) or no treatment (blue). Mean ± SD (n = 2 technical replicates). The averaged expression level of *pHes7-luc* iPSM colonies without dox was set to 1. Two-tailed t-test, *P < 0.05, **P < 0.01. (**F**) The premature expression of CLOCK/BMAL1 in the iPSM and gastruloids interfered with the segmentation clock oscillation and somitogenesis-like process.

## Discussion

Our present study showed that premature expression of circadian key components CLOCK/BMAL1 severely interferes with the ultradian rhythm of the segmentation clock in iPSM and gastruloids.

We have previously reported that during the early to mid-developmental stage, there are multiple molecular mechanisms that underlie the strict suppression of circadian TTFLs, such as the post-transcriptional suppression of CLOCK protein (17, 18) and the exclusive cytoplasmic localization of PER proteins (16). Furthermore, we have also reported that the maternal circadian clock cannot entrain the fetus until the establishment of the fetal circadian clock itself (17). These results suggest that the circadian rhythm in mammalian embryos is rigorously suppressed by the multilayered inhibitory mechanisms during the early to mid-developmental stage. During the multilayered suppression of circadian clock oscillation, the ultradian temporal oscillation of *Hes7* expression, segmentation clock, proceeds and forms the spatial repetitive structure of somites.

In the present study, we investigated the effect of the CLOCK/BMAL1-mediated activation of *Per1* transcription on the segmentation clock oscillation by using the iPSM differentiated from ESCs. It was suggested that, similar to the undifferentiated ESCs, circadian clock oscillation is suppressed in the iPSM by the common mechanisms to the ESCs and early embryos (**see Fig. S2**). Recently, it was reported that hundreds of genes including *Per1* also oscillates in the same phase as *Hes7* ultradian rhythm in *in vitro*-PSM of both mouse and humans (30), suggesting that *Per1* is deviated and free from the circadian gene regulatory mechanism of TTFL. These findings are consistent with the previously reported observations indicating that the multilayered inhibitory mechanisms including post-transcriptional inhibition of CLOCK and the predominant cytoplasmic accumulation of PER1 do not allow the oscillation of circadian TTFL (17, 18). Interestingly, although the expression of BMAL1 protein was observed even in ESCs (17), the dox-induced CLOCK sole expression in ESCs resulted in the only partial upregulation of E-box driven circadian clock genes (**Fig. S3**), raising the possibility that the endogenously expressed BMAL1 might be post-translationally modified to not function. Therefore, in this study, we used ESC lines carrying both the dox-inducible *Clock* and *Bmal1* genes as a model system of premature expression of CLOCK/BMAL1 (**see Fig. 1C**).

We demonstrated that the expression of CLOCK/BMAL1 affected the WNT, MAPK, and NOTCH signaling pathways related to *Hes7* oscillation in iPSM (**see Fig. 5 A and B**). In addition, the premature expression of CLOCK/BMAL1 resulted in not only the up-regulation of *Per1* expression but also the expressions of *Hes7, Aloxe3*, and *Vamp2*, localized adjacently on the *Per1* genomic locus (**see Fig. 2 A and B, Fig. 5 C–E, Fig. S6 C–E**). In the iPSM, the up-regulation of these gene expressions has already been induced after the 2-hour dox treatment (**see Fig. 2 A and B, Fig. 5 C–E**). Considering that the *Per1* promoter harbors E-box elements with which CLOCK/BMAL1 heterodimer has a much higher affinity than the other genomic region (31), the immediate up-regulation of genes near *Per1* gene locus after the induction of CLOCK/BMAL1 expressions could be caused by the ripple effect (32). On the other hand, the bioluminescence from *Hes7*-promoter driven luciferase reporters in the iPSMs not only lost cycling but also decreased signal intensity in approximately 2 hours after the dox addition (**see Fig. 3D**), indicating that the expression of CLOCK/BMAL1 in the iPSM has also inhibitory effects on the *Hes7* gene expressions. Among components involved in the *Hes7*-regulatory signaling pathways, expression of CLOCK/BMAL1 induced some negative regulators, such as the *Dusp* phosphatase family (33) in the MAPK signaling pathway, *Sfrp* in the WNT signaling pathway (34), and *Lfng* in the NOTCH signaling pathway (35) (**see Fig. 5B**). Therefore, the premature expression of CLOCK/BMAL1 first may upregulate *Hes7* transcription and induce subsequent downregulation of *Hes7* gene expression by the induction of the negative regulators in addition to the HES7 autoinhibition. Consequently, the premature expression of the circadian components CLOCK/BMAL1 interfered with *Hes7* oscillation by perturbing the *Hes7* expression through various pathways.

In this study, we used a mouse embryonic organoid, gastruloids, as an *in vitro* recapitulation model of somitogenesis-like process (27). The premature expression of CLOCK/BMAL1 in the gastruloids disrupted not only the *Hes7* oscillation but also the striped structure of the somite marker, *Uncx4*.*1* (**see Fig. 4M**). Because the RNA-seq analysis data showed that hundreds of genes were affected by the induction of CLOCK/BMAL1 (**see Fig. 5A**), the possibility cannot be denied that the premature expression of CLOCK/BMAL1 affects cell fates or characters. However, the posterior structure in the gastruloids was held even after the induction of CLOCK/BMAL1 and then continued to extend, concomitant with the decrease of *Hes7* bioluminescence signals and the arrest of the *Hes7* oscillation (**see Fig. 4K**), suggesting that the premature expression of CLOCK/BMAL1 interfered with the somitogenesis process by perturbing *Hes7* oscillation of the segmentation clock.

In vitro recapitulation of embryonic process using iPSM and gastruloids has differences such as no brain tissues comparing with in vivo process. However, key regulators of somitogenesis we focus on in this study are expressed similarly between embryos and gastruloids using single-cell RNA sequencing and spatial transcriptomics (27), and the in vitro recapitulation model enables to analyze the *Hes7* oscillation in more detail using real-time imaging without maternal effects.

Our findings shown in this study indicated that the CLOCK/BMAL1, key components regulating the circadian TTFL, affected and interfered with the segmentation clock. Considering that transcriptional activation of CLOCK/BMAL1 is essential for the circadian regulatory networks, these results suggest that the strict suppression of circadian molecular oscillatory mechanisms during the early stage embryos is inevitable for the intact developmental process in mammals. Therefore, this may be the biological and physiological significance of the delayed emergence of circadian clock oscillation and the rhythm conversion observed in mammalian development.

## Materials and Methods

### Cell culture

KY1.1 ESCs (7), referred to as ESC in the text, and *Per2*^*Luc*^ ESCs (5, 36) were maintained as described previously (17). E14TG2a ESCs carrying *Hes7*-promoter-driven luciferase reporters (25), referred to as *pHes7-luc* ESCs in the text, were maintained without feeder cells in DMEM (Nacalai) supplemented with 15% fetal bovine serum (Hyclone), 2 mM L-glutamine (Nacalai), 1 mM nonessential amino acids (Nacalai), 100 µM StemSure^®^ 2-mercaptoethanol solution (Wako), 1 mM sodium pyruvate (Nacalai), 100 units/mL penicillin and streptomycin (Nacalai), 1000 units/mL leukemia inhibitory factor (Wako), 3 µM CHIRON99021 (Wako or Tocris Biosciences), and 1 µM PD0325901 (Wako) with 5% CO_2_ at 37°C.

### Transfection and establishment of cell lines

ESCs stably expressing dox-inducible *Clock/Bmal1* or *Clock/Bmal1DN* (I584X) were established as described previously (17). For *TetO-Clock/Bmal1* or *TetO-Clock/Bmal1DN* ESCs, KY1.1 ESCs or *pHes7-luc* ESCs were transfected using 10.5 µl of FuGENE 6 mixed with 1 µg of pCAG-PBase, 1 µg of PB-TET-Clock (17), 1 µg of PB-TET-Bmal1 or PB-TET-Bmal1DN (I584X), 1 µg of PB-CAG-rtTA Adv, and 0.5 µg of puromycin selection vector. The transfected cells were grown in a culture medium supplemented with 2 µg/mL puromycin for two days. The ESC colonies were picked and checked by qPCR after treatment with 500 ng/mL dox. For PB-TET-Bmal1 and PB-TET-Bmal1DN (I584X), Bmal1 cDNA and Bmal1DN (I584X) cDNA (26) were cloned into a PB-TET vector (37). For the *TetO-Clock Per2*^*Luc*^ ESCs, *Per2*^*Luc*^ ESCs were established as described previously (17).

### Bioluminescence imaging

The iPSM colonies were differentiated from the *pHes7-luc* ESCs and *Per2*^*Luc*^ ESCs as described previously (25). The *Per2*^*Luc*^ ESCs were cultured without feeder cells in the ES medium containing 3 µM CHIRON99021 and 1 µM PD0325901 before in vitro differentiation. Bioluminescence imaging of single *pHes7-luc* iPSM colonies and *Per2*^*Luc*^ iPSM colonies was performed in gelatin-coated 24-well black plates or 35-mm dishes (26). DMEM was used that was supplemented with 15% Knock-out Serum Replacement (KSR), 2 mM L-glutamine, 1 mM nonessential amino acids, 1 mM sodium pyruvate, 100 units/mL penicillin and streptomycin, 0.5% DMSO, 1 µM CHIRON99021, and 0.1 µM LDN193189 (Sigma) containing 1 mM luciferin and 10 mM HEPES. For live imaging of single iPSM colonies using an EM-CCD camera, each iPSM colony was cultured on a fibronectin-coated glass base dish for 6 h, and images were acquired every 5 min with an exposure time of 10 sec (control) or 2.5 sec (*Clock/Bmal1* induction) under 5% CO_2_ using an LV200 Bioluminescence Imaging System (Olympus).

Gastruloids were generated as described in a previous report (27). In total, 200–250 live cells were plated in 40 µl of N2B27 medium into each well of a U-bottomed nontissue culture-treated 96-well plate (Greinier 650185). After a 96-hr cultivation, the gastruloids were embedded in 10% Matrigel (Corning 356231) containing 1 mM luciferin. For live imaging of single gastruloids, the images were acquired every 5 min with an exposure time of 3.5 sec (*Clock/Bmal1* induction) or 10 sec (control) under 5% CO_2_ using the LV200 system. The Videos were analyzed using the ImageJ software (38). Kymographs of the averaged bioluminescence intensity along the straight or segmented line of 5-pixel width were generated using the plug-in KymoResliceWide.

### *In situ* hybridization

Hybridization chain reaction (HCR) v3 was performed as described previously (27, 39) using reagents procured from Molecular Instruments. *Uncx4*.*1* HCR probe (Accession NM_013702.3, hairpin B1) was labeled with Alexa Fluor 488.

### Quantitative RT-PCR

The iPSM colonies, ESCs, and gastruloids were washed with ice-cold PBS, and total RNA was extracted using Isogen reagent (Nippon Gene) or miRNeasy Mini Kits (QIAGEN) according to the manufacturer’s instructions. To remove the feeder cells from ESCs cultured on a feeder layer, the cells were treated with trypsin, and then the mixed cell populations were seeded on gelatin-coated dishes and incubated for 25 min at 37°C three times in ES cell medium. Non-attached ESCs were seeded in a gelatin-coated dish overnight and then treated with or without 500 ng/mL doxycycline for 6 hr. The iPSM colonies and gastruloids were treated with or without 1000 ng/mL doxycycline for 2 hr. First-strand cDNAs were synthesized with 1000 or 280 ng of total RNA using M-MLV reverse transcriptase (Invitrogen) according to the manufacturer’s instructions. Quantitative PCR analysis was performed using the StepOnePlus™ Real-Time PCR system (Applied Biosystems) and iTaq™ Universal SYBR Green Supermix (Bio-Rad Laboratories). Standard PCR amplification protocols were applied, followed by dissociation-curve analysis to confirm specificity. Transcription levels were normalized to the level of β-actin. The following primer sequences were used:

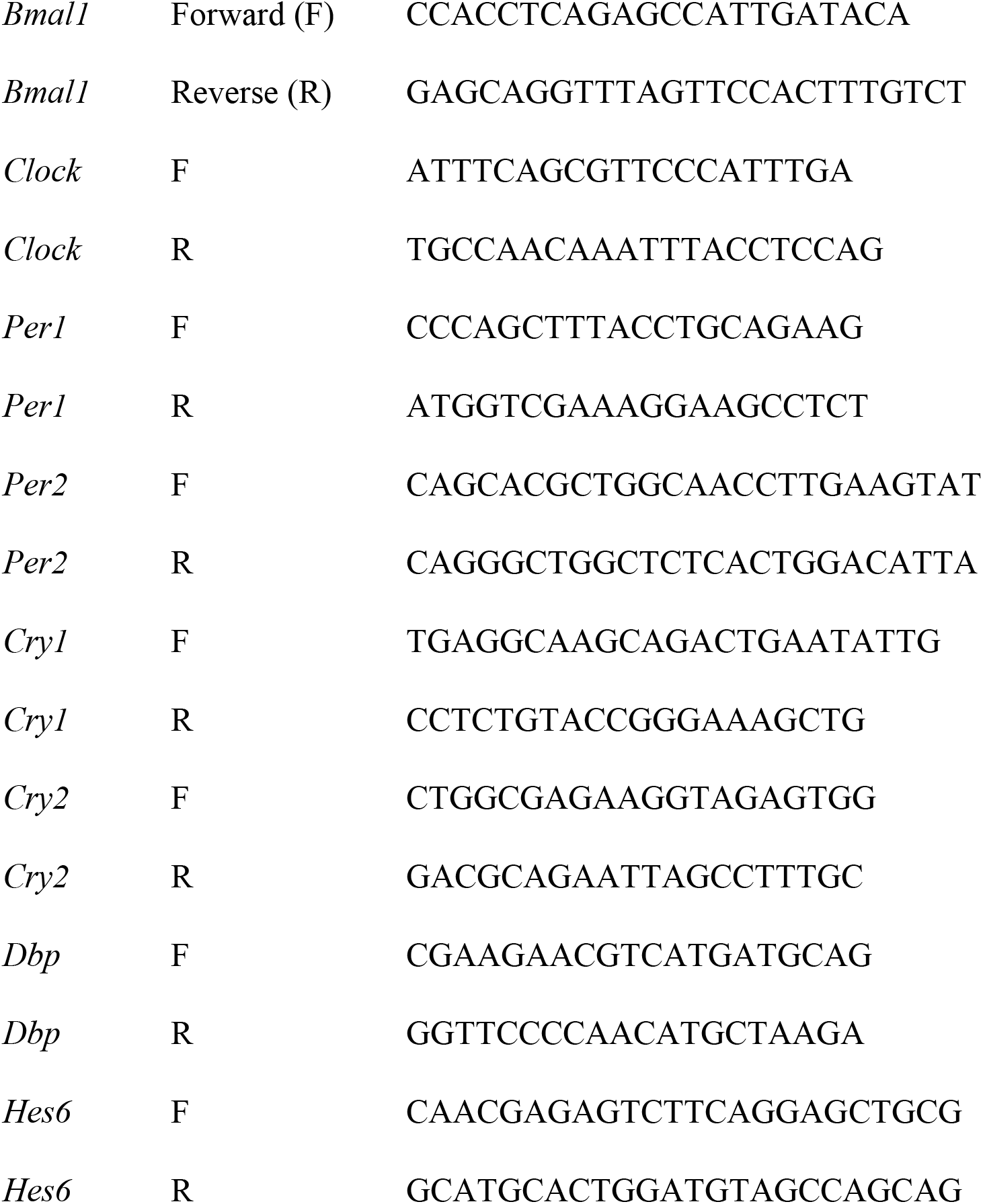
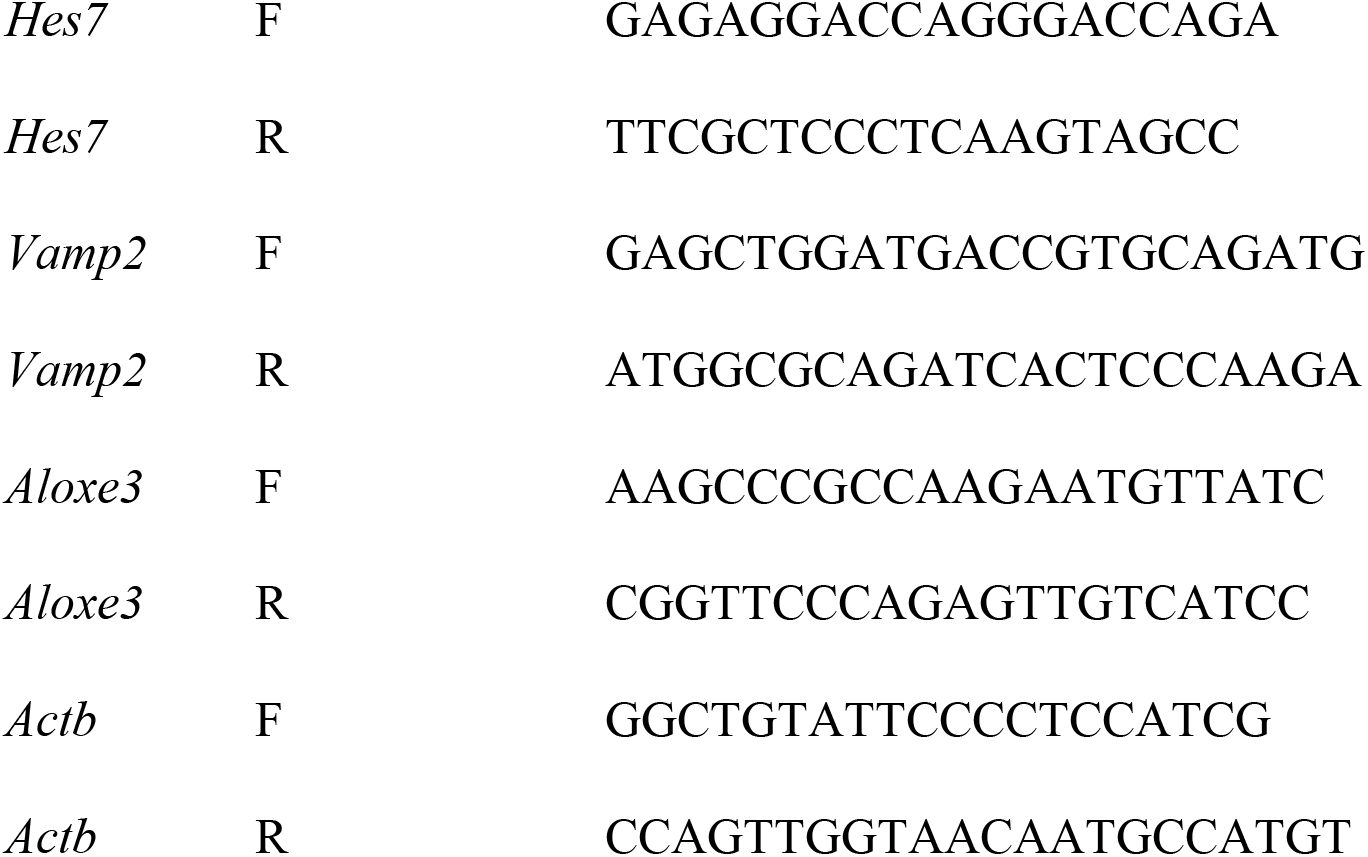

### RNA-seq

The iPSM colonies and ESCs were washed with ice-cold PBS, and total RNA was extracted using miRNeasy Mini Kits (QIAGEN) according to the manufacturer’s instructions. Total RNA sequencing was conducted by Macrogen Japan on an Illumina NovaSeq 6000 with 101-bp paired-end reads. After trimming the adaptor sequences using Trimmomatic (40), the reads that mapped to ribosomal DNA (GenBank: BK000964.1) (41) were filtered out and the sequence reads were mapped to the mouse genome (GRCm38/mm10) using STAR (42), as described previously (16). To obtain reliable alignments, reads with a mapping quality of less than ten were removed using SAM tools (43). The known canonical genes from GENCODE VM23 (44) were used for annotation, and the reads mapped to the gene bodies were quantified using Homer (45). The longest transcript for each gene was used for gene-level analysis. We assumed that a gene was expressed when there were more than 20 reads mapped on average to the gene body. Differential gene expression in the RNA-seq data was determined using DESeq2 with thresholds of FDR < 0.05, fold change > 1.5, and expression level cutoff > 0.1 FPKM (46). WebGestalt was used for KEGG pathway enrichment analysis (47). In the RNA-seq data using iPSM colonies, the reads mapped in the promoter (chr11:69115096-69120473) and 3′UTR (chr11:69122995-69123324) of *Hes7* were filtered out to eliminate transcripts from the *pHes7-luc* reporter transgene. The heatmaps of gene expression and KEGG pathways were generated with R using the pheatmap and pathview packages, respectively.

### Immunostaining

The iPSM colonies were fixed in cold methanol for 15 min at room temperature. The fixed iPSM was blocked with 1% BSA or 5% skim milk overnight at 4°C and then incubated with anti-CLOCK mouse antibody (CLSP4) (48), anti-BMAL1 mouse antibody (MBL, JAPAN), anti-BMAL1 guinea pig antibody (16), or anti-PER1 rabbit antibody (AB2201, Millipore) overnight at 4°C. After washing in 1% BSA, the iPSM colonies were incubated with a CF^™^488A-conjugated donkey anti-mouse IgG (Nacalai), Cy3-conjugated goat anti-guinea pig IgG (Jackson), DyLight^™^488-conjugated donkey anti-rabbit IgG (Jackson) for 2 hr at 4°C, and the nuclei were stained with TO-PRO®-3 1:1000 (Thermo Fisher Scientific, USA) for 10–20 min. The iPSM colonies were washed in 1% BSA and observed using an LSM510 or 900 confocal laser scanning microscope (Zeiss).

## Data availability

RNA sequence data are available at the Gene Expression Omnibus. All other datasets generated in this study are available from the corresponding author upon reasonable request.

## Acknowledgments

We thank the Yagita lab members for technical assistance. This work was supported in part by grants-in-aid for scientific research from the Japan Society for the Promotion of Science to Y.U. (19K06679) and K.Y. (18H02600), the Cooperative Research Program (Joint Usage/Research Center program) of the Institute for Frontier Life and Medical Sciences, Kyoto University (K.Y. and G.K.)

## Supporting information

**Fig. S1.**
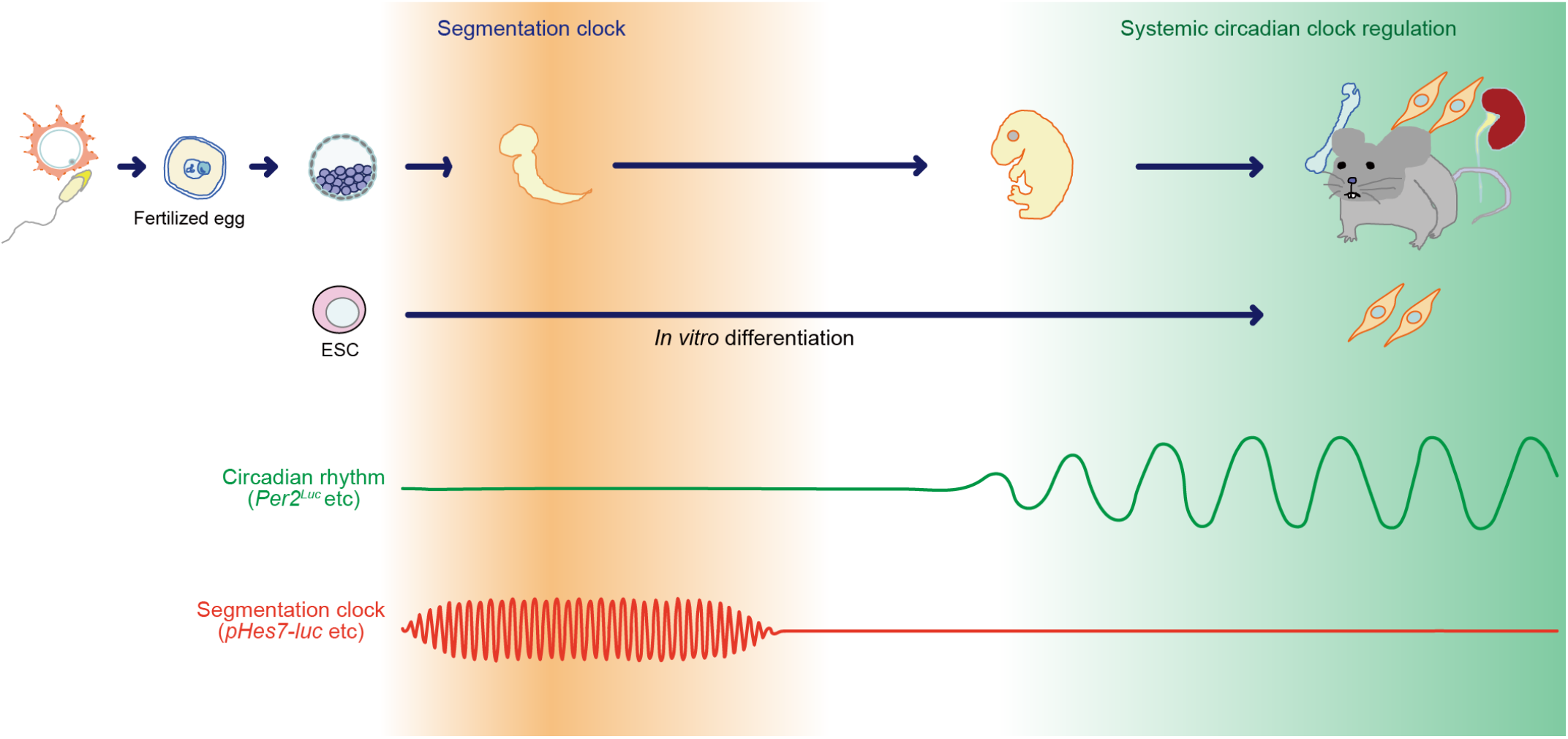
Mutually exclusive appearance of segmentation clock and circadian clock. In mammals, two different types of rhythm appear sequentially during the developmental process. One is the ultradian rhythm by segmentation clock, which controls somitogenesis. The other one is the circadian oscillation, which regulates the predictive adaptation of physiological functions to the day–night environmental cycle.

**Fig. S2.**
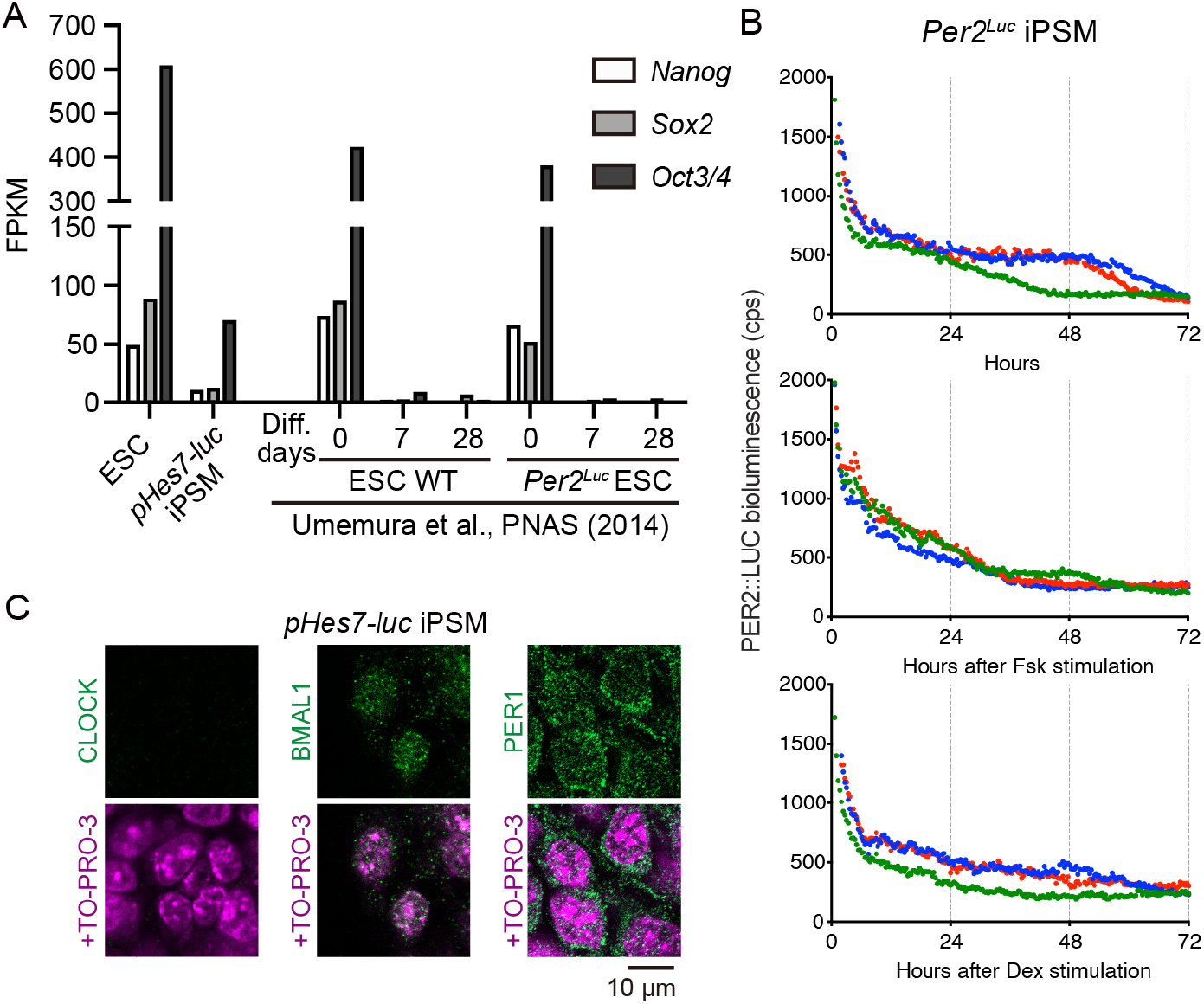
The iPSM has no apparent circadian clock oscillation. **(A)** The expression levels of pluripotent marker genes, *Nanog, Sox2*, and *Oct3/4*, were measured using RNA-seq in ESCs and *pHes7-luc* iPSM. The data were shown with *in vitro* 0, 7-, and 28-day differentiated WT ESCs and *Per2*^*Luc*^ ESCs (GSE61184). No circadian clock oscillates in the *in vitro* 7-day differentiated ESCs and the circadian clock oscillation starts to emerge after 14 days of differentiation culture (16, 17). **(B)** Bioluminescence traces of the iPSM differentiated from *Per2*^*Luc*^ ESCs. The iPSM was stimulated with 10 µM forskolin (Fsk, middle) or 100 nM dexamethasone (Dex, bottom). n = 3 biological replicates. **(C)** Representative immunostaining of CLOCK, BMAL1, and PER1 in the iPSM. The immunostaining represented the suppression of CLOCK proteins in contrast to the expression of BMAL1. Furthermore, although the nuclei indicated the quite faint signals, PER1 protein in the iPSM is still predominantly accumulated in the cytoplasm.

**Fig. S3.**
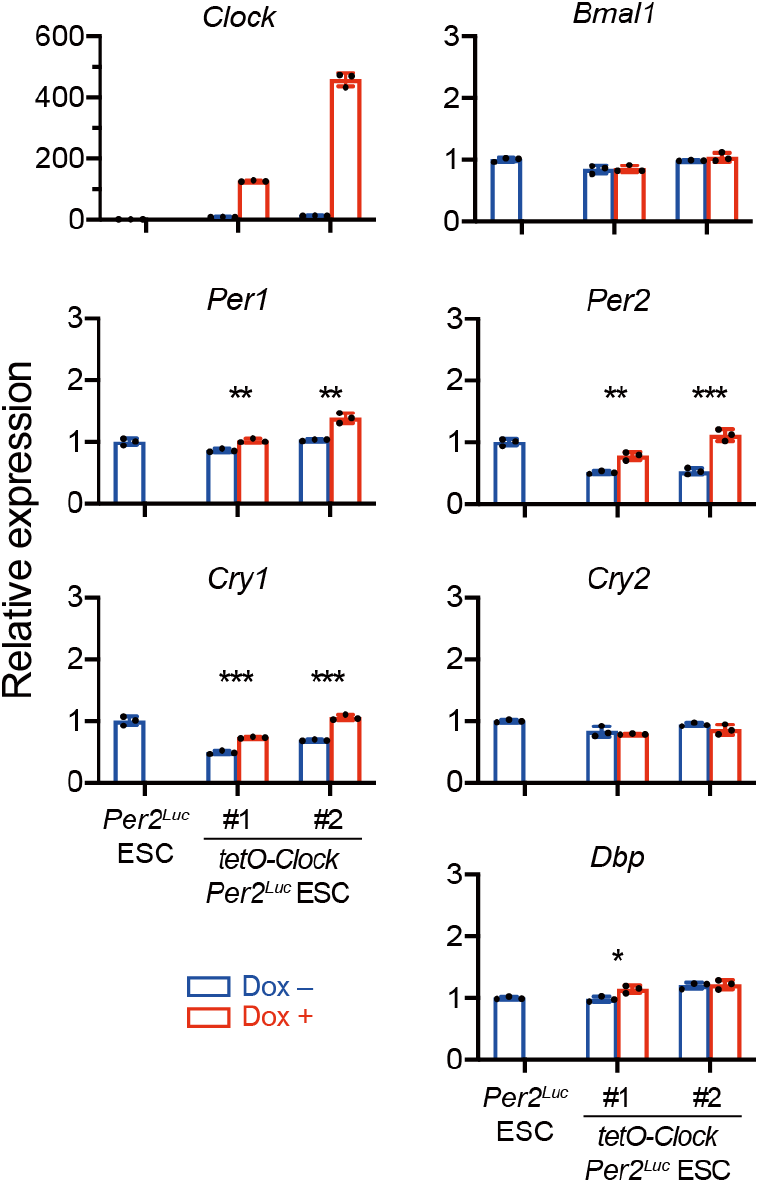
The core circadian clock gene expressions in the ESCs harboring dox-inducible *Clock*. Although the expression of BMAL1 protein was observed even in ESCs (17), dox-dependent sole expression of *Clock* was insufficient for the E-box-driven expression of clock genes such as *Per1/2* and *Cry1/2* in undifferentiated ESCs, raising the possibility that the endogenously expressed BMAL1 might be post-translationally modified to not function. Colored boxes indicate 500 ng/mL dox treatment for 6 hr (red) or no treatment (blue). Each number indicates clone number. Mean ± SD (n = 3 biological replicates). The averaged expression level of *Per2*^*Luc*^ ESCs without dox was set to 1. Two-tailed t-test, *P < 0.05, **P < 0.01, ***P < 0.001.

**Fig. S4.**
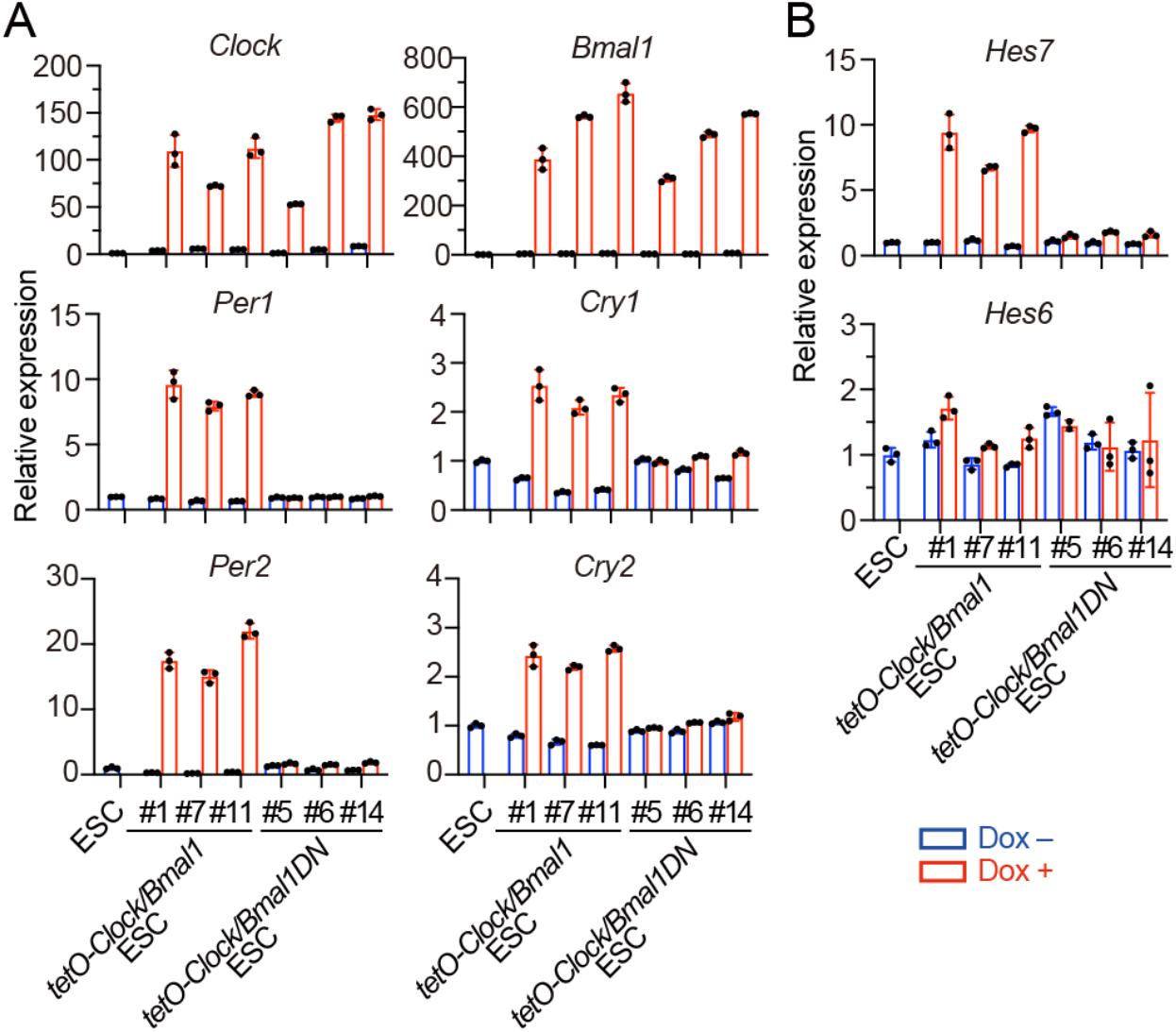
*Clock/Bmal1* gene expressions in ESCs upregulated not only circadian clock genes but also *Hes7* gene expression. (**A, B**) qPCR of core circadian clock gene (**A**) and *Hes*7 or *Hes6* gene (**B**) expression in ESCs harboring dox-inducible *Clock* and *Bmal1* or *Clock* and *Bmal1DN*. Colored boxes indicate the presence (red) or absence (blue) of 500 ng/mL dox treatment for 6 hr. Each number indicates clone number. Mean ± SD (n = 2–3 biological replicates). The average expression level of ESCs without dox was set to 1.

**Fig. S5.**
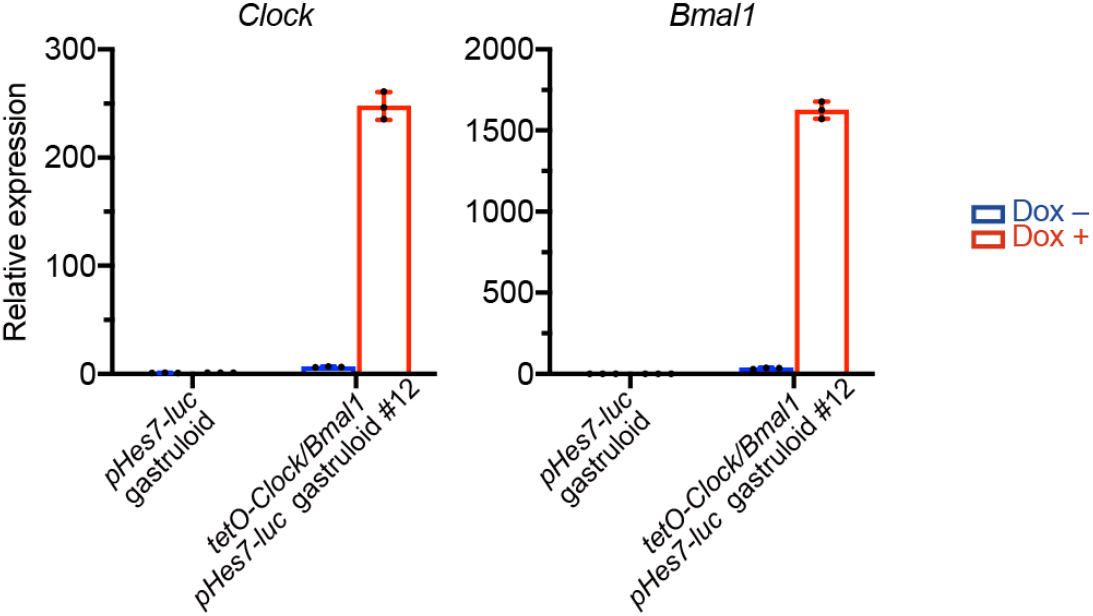
qPCR of *Clock* and *Bmal1* mRNA in the indicated gastruloids. Colored boxes indicate 1000 ng/mL dox treatment for 2 hr (red) or no treatment (blue). Each number indicates clone number. Mean ± SD (n = 3 biological replicates). The average expression level of *pHes7-luc* gastruloids without dox was set to 1.

**Fig. S6.**
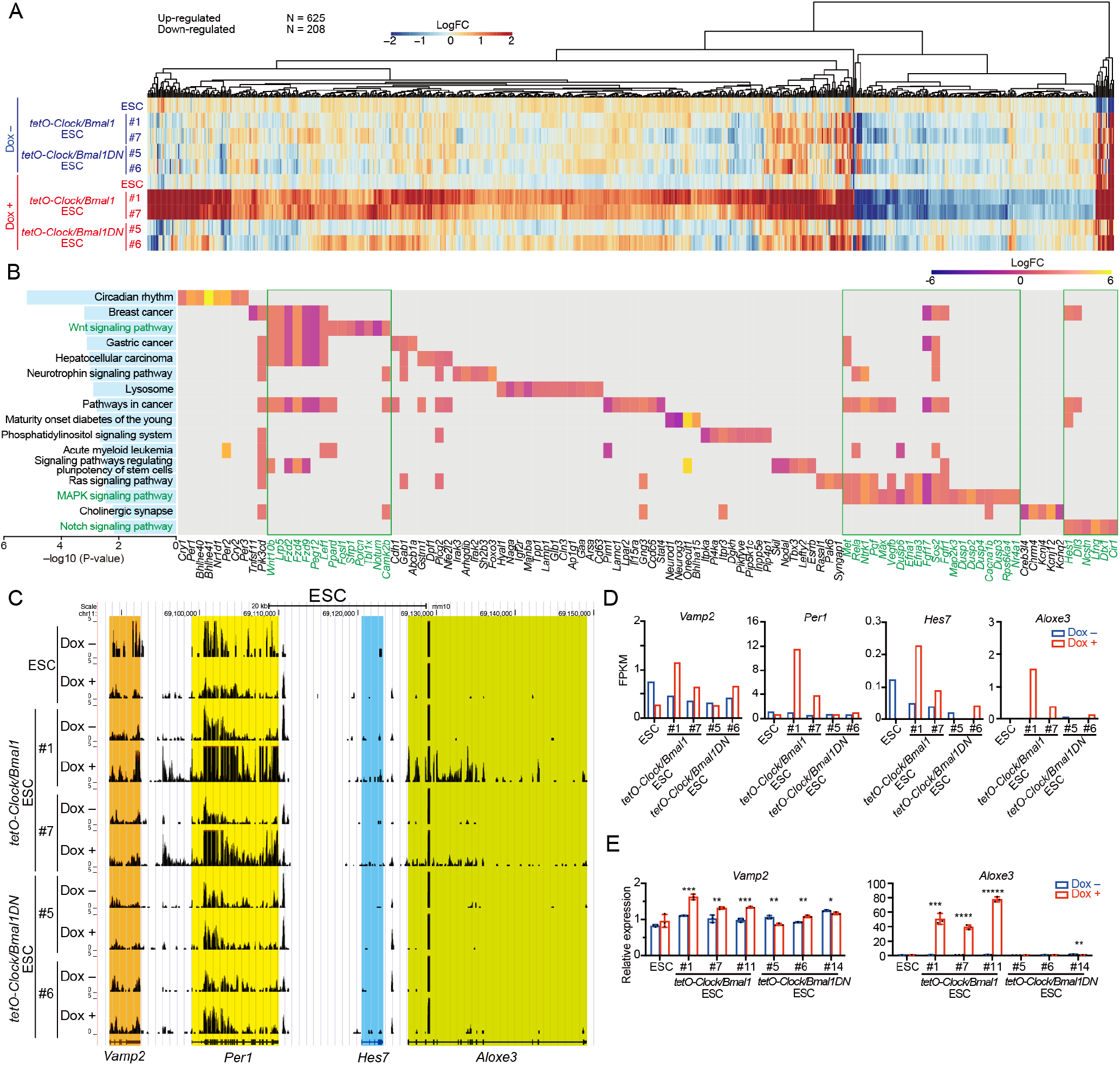
*Clock/Bmal1* gene expressions in ESCs affected the *Hes7*-related signaling pathways and upregulated the expression of the contiguous genes *Vamp2, Per1, Hes7*, and *Aloxe3*. (**A**) Upregulated and downregulated differentially expressed genes in the indicated ESCs treated with dox (FDR < 0.05, FC > 1.5). (**B**) KEGG pathway analysis of the DEGs. Each pathway was indicated with each transformed p-value. The ranked pathways contained several common genes in the WNT, MAPK, and NOTCH signaling pathways. (**C**) UCSC genome browser views of RNA-seq data of contiguous genes, including *Vamp2, Per1, Hes7*, and *Aloxe3*. The reads shown are normalized average reads per 10 million total reads in 10-bp bins. (**D**) mRNA expression of *Vamp2, Per1, Hes7*, and *Aloxe3* in the indicated ESCs measured using RNA-seq. (**E**) Validation of *Vamp2* and *Aloxe3* in the indicated ESCs by qPCR. Colored boxes indicate the use of 500 ng/mL dox treatment for 6 hr (red) or untreated cells (blue). Each number indicates clone number. Mean ± SD (n = 3 biological replicates). Two-tailed t-test, *P < 0.05, **P < 0.01, ***P < 0.001, ****P < 0.0001, *****P < 0.00001.

